# Warming indirectly increases invasion success in food webs

**DOI:** 10.1101/2020.07.20.211516

**Authors:** Arnaud Sentis, Jose M. Montoya, Miguel Lurgi

## Abstract

Climate warming and biological invasions are key drivers of biodiversity change. Their combined effects on ecological communities remain largely unexplored. We investigated the direct and indirect influences of warming on invasion success, and their synergistic effects on community structure and dynamics. Using size-structured food web models, we found that warming increased invasion success. The direct physiological effects of warming on invasions were minimal in comparison to indirect effects mediated by changes on food web structure and stability. Warmed communities with less connectivity, shortened food chains and reduced temporal variability were more susceptible to invasions. The directionality and magnitude of invasions effects on food webs varied across warming regimes. Warmer communities became smaller, more connected, and with more predator species when invaded than their colder counterparts. They were also less stable and their species more abundant. Considering food web structure is crucial to predict invasion success and its impacts under warming.

## INTRODUCTION

Climate warming and biological invasions constitute two of the most pervasive drivers of global change (Nelson 2005; Díaz *et al.* 2020). Both drivers strongly impact ecosystems, causing not only species loss, but also affecting ecological interactions and the structure of interaction networks (Stachowicz *et al.* 2002; Holzapfel & Vinebrooke 2005; Romanuk *et al.* 2009; Britton *et al.* 2010; Huang *et al.* 2011; Lurgi *et al.* 2012b; Lu *et al.* 2013; Lurgi *et al.* 2014; Zhang *et al.* 2017). Climate warming and species invasions can act synergistically on ecosystems due to, for example, impacts of climate change on species niche range dynamics (Thuiller *et al.* 2008; Elith & Leathwick 2009; Tylianakis & Morris 2017), which differentially influence species’ ability to colonise new habitats and thus invade new communities. Species range shifts affect not only species composition, but also the structure of species interaction networks creating novel communities. For example, by promoting species range shifts, climate warming can trigger the loss of specialised interactions and changes the body size ratio between predator and prey species, which in turn can influence predator control on prey populations (Lurgi *et al.* 2012b, a). Yet, we know surprisingly little about how invasions and climate change act together to affect species and links in ecosystems.

Previous studies have shown that warming can enhance invasions by increasing survival and reproduction of introduced species (Mandrak 1989; Johnson & Evans 1990; Stachowicz *et al.* 2002; Logan *et al.* 2003; Britton *et al.* 2010; Huang *et al.* 2011; Ricciardi *et al.* 2017). However, warming can also lead to the opposite effect by decreasing the potential for invaders to occupy new niches (Bradley *et al.* 2010; Bertelsmeier *et al.* 2013). Recent evidence suggests that warming influence on invasion success may depend on how warming influences trophic interaction strength and the persistence of native predators or competitors (Holzapfel & Vinebrooke 2005; Lu *et al.* 2013; Seifert *et al.* 2015). On the one hand, warming can prevent invasions by increasing top-down control on the invader prey (Bradley *et al.* 2010; Lu *et al.* 2013; Lu *et al.* 2016). On the other hand, warming can enhance invasions by releasing top-down control following predator extinctions (Holzapfel & Vinebrooke 2005). Overall, previous studies reported various outcomes on the effects of warming on invasions. Our limited understanding of their causes poses the challenge for gaining a better understanding of the indirect effects of warming on species and communities. Indirect effects of warming on community structure and species interactions are often stronger than its direct effects on physiology and demography (Ockendon *et al.* 2014). This suggests that investigating warming effects on communities and the complex networks of interactions that structure them is a first step to address this challenge.

Ecologists have developed mechanistic frameworks to identify key processes underlying temperature effects on trophic interactions and networks (Binzer *et al.* 2012; Burnside *et al.* 2014; Fussmann *et al.* 2014; Gilbert *et al.* 2014; Sentis *et al.* 2014). One first important finding is that, since consumer metabolic rates often increase faster with temperature than their feeding rates, most consumers become less efficient at processing matter and energy at warmer temperatures (Vucic-Pestic *et al.* 2011; Fussmann *et al.* 2014; Iles 2014). This reduction of energetic efficiency lessens energy flow between trophic levels and, if resulting in weakened interaction strengths, it stabilizes food-web dynamics by reducing population fluctuations (Rip & McCann 2011; Binzer *et al.* 2012; Gilbert *et al.* 2014). A second important finding is that elevated temperatures increase consumer extinction risk when metabolic demands exceed ingestion rates, leading to consumer starvation and extinction (Petchey *et al.* 1999; Rall *et al.* 2010; Sentis *et al.* 2020). Whether these changes would favour invasion success is unclear, as previous studies exploring the role of community structure and dynamics in preventing or facilitating invasions success have not considered modifications in communities driven by climate change (e.g. Romanuk *et al.* 2009; Lurgi *et al.* 2014).

In parallel to studies focusing on the effects of warming, much effort has been devoted to understanding how invasions impact ecosystems (Hui & Richardson 2019). Several models have unveiled the role of food web structure such as species richness, complexity or the heterogeneity of distribution of interactions in preventing invasions. These models have also suggested that invasions in food webs tend to decrease species richness and shorten food chains (e.g. Romanuk et al. 2009; Lurgi et al. 2014). However, a more recent theoretical investigation suggests the opposite, with invasions being instead beneficial for maintaining species richness and ecological functions (Zhang et al. (2017). We need comprehensive mechanistic frameworks incorporating both warming and invasion and their effects on complex communities to better understand and predict their synergistic effects.

Here, we explored the combined effects of warming and invasions on food webs using a theoretical model. Given our current understanding of the effects of warming on natural communities on the one hand (Binzer *et al.* 2012; Sentis *et al.* 2017; Boukal *et al.* 2019), and of the invasion process in complex food webs on the other (Romanuk et al. 2009; Lurgi et al. 2014; Hui & Richardson 2019), along with previous studies showing a positive influence of weakened top-down control on invasions (Holzapfel & Vinebrooke 2005), we hypothesise that warming increases invasion success if it decreases top-down control or causes predator extinctions. We further hypothesise warming to increase connectivity and shorten food chains in response to species loss at higher trophic levels. These effects of warming should in turn destabilise community dynamics (Boukal *et al.* 2019).

To test these hypotheses, we investigated how temperature can influence invasions on complex food webs comprised of 30 species. Our model simulates population dynamics in food webs following bio-energetic principles of species life histories and interactions. The model incorporates the temperature dependency of biological rates allowing the exploration of a wide range of temperature regimes. We aim at gaining a better understanding of (1) warming effects on invasion success in food webs and (2) the ecological consequences of invasions on food web structure in warmed communities.

## MATERIAL AND METHODS

We modelled community dynamics in complex food webs using a size-structured bio-energetic community model consisting of a set of ordinary differential equations (ODEs) incorporating the effects of species growth and ecological interactions (Yodzis & Innes 1992; Brose *et al.* 2006). Food webs were generated using the niche model (Williams and Martinez 2000). The effect of temperature on population dynamics was incorporated into ODEs by introducing thermal dependencies of relevant model parameters. A series of numerical simulations were then computed and species invasions were modelled as the addition of a new species into the community (Lurgi et al. 2014). Simulation results were analysed to assess the effects of temperature on (1) food web proprieties (structure, stability and total biomass) before invasion, invasion success, and (3) the effects of invasions on community structure and stability.

### Food web generation

Food webs were generated using the niche model (Williams & Martinez 2000). With only two parameters (number of species (*S*) and network connectance (*C*), i.e. the fraction of links out of all possible ones) this model generates networks that resemble real food web structure (Williams & Martinez 2000). We generated food webs comprising 30 species and with 10% connectance. These values are within the ranges reported for empirical food webs (Williams & Martinez 2000). We kept these values fixed across experiments to avoid the confounding effects of variation in species richness and connectance.

### Non-linear model for population dynamics

To simulate network dynamics of species populations biomass we used an allometric bio-energetic model adapted from its original formulation by Yodzis and Innes (1992). This model defines interactions strengths between prey and predator according to their body mass ratios (Brose *et al.* 2006) and has been used to investigate warming and invasion effects on complex food webs (Binzer *et al.* 2011; Lurgi *et al.* 2014). Eqn. 1 gives dynamics of basal resource species. These grow logistically with an intrinsic growth rate *r*_*i*_ and a carrying capacity *K*_*i*_. Consumers gain biomass according to Eqn. 2:

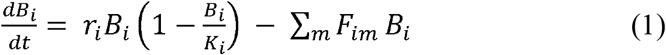

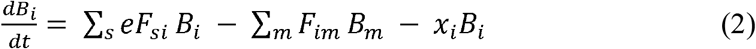

where *B*_*i*_ is the biomass of species *i*; *e* is the assimilation efficiency of predators when ingesting prey (kept constant across consumer-resource species pairs at a value *e* = 0.85 for carnivorous species (following Yodzis & Innes 1992)); *x*_*i*_ is the metabolic rate at which biomass of consumers is lost from the system due to respiration and other metabolic processes. *F*_*ij*_ is a function that describes the feeding relationship between prey *i* and predator *j* and is defined by the functional response:

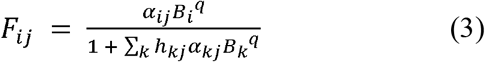

where α_*ij*_’s are the elements of a quantitative version of the adjacency matrix *A*, describing the food web obtained according to the procedure explained above (*Food web generation*), and that represent the attack rates of predator species *j* on prey species *i* (Eqn. 5a). *h*_*ij*_ is the handling time, i.e. the average time spent by an individual of predator species *i* handling and digesting an individual of prey species *i*. The shape of the functional response curve is controlled by the parameter *q* (i.e. the Hill exponent). We kept *q* constant across interacting species pairs at a value of 1.2 to simulate an intermediate response type between Type II (hyperbolic, *q* = 1) and Type III (sigmoidal, *q* = 2), as in (Binzer et al. 2011, 2016).

Growth, metabolic, attack rates, and handling times are functions of species body masses and temperature. Body mass of species *i* scales according to its position in the food web:

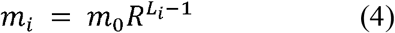

where *m*_*0*_ is the body size of basal species in the food web and set here to *m*_*0*_ = 0.01g, *R* is the average predator-prey body mass ratio of all trophic interactions in the food web and was set here to *R* = 10^2^, *L*_*i*_ is the prey-averaged trophic level of species *i* (Williams & Martinez 2004). Allometric and thermal dependencies of model parameters were incorporated as follows (Binzer et al. 2016):

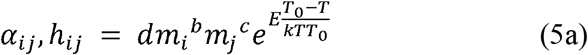

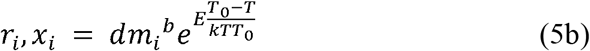

where *d* = *e*^*I*^ is a rate-specific constant calculated for a species with body mass of 1g and at a reference temperature *T*_0_ = 20°C (293.15K), *m*_i_ and *m*_j_ are the body masses of species *i* and *j*, respectively, *b* and *c* are rate-specific allometric exponents. The temperature dependence term is a version of the Arrhenius equation in which *E* is the rate-specific activation energy and *k* is the Boltzmann constant. *T* is the current temperature of the system in Kelvin. *T*_0_ is the reference temperature at which the rate value is equal to the rate-specific constant *d*. Values and units for the parameters in Eqns. 5a and 5b are presented in Table 1.

Species carrying capacity was assumed to be independent of temperature since empirical evidence for the thermal dependency of carrying capacity is inconclusive (Fussmann *et al.* 2014; Gilbert *et al.* 2014; Uszko *et al.* 2017; Bernhardt *et al.* 2018). Furthermore, we wanted to avoid biases in the invasion experiments due to the intrinsic limit to community biomass caused by the negative temperature dependence of carrying capacity, which would, in turn, influence invasion success. We thus focused on the effects of temperature on species life-history traits such as reproduction, death and species interactions (i.e. attack rates and handling times) and not on the maximum population density of the basal resources.

**Table 1.** Parameter values for mass and temperature dependencies of r, α, h and x. Parameters are in biomass units, i.e. per unit of mass of predator or prey. Parameters extracted / calculated from: growth (*r* in [1/s], (Savage et al., 2004)), metabolism (*x* in [1/s], (Ehnes et al., 2011)), attack rate (a in [m^2^ s^−1^]) and handling time (*h* in [s]), both calculated from (Rall et al., 2012). Metabolic rates were calculated using the conversion factor from Peters (1983).

| | $r_i$ | $\alpha_{ij}$ | $h_{ij}$ | $x_i$ |
| --- | --- | --- | --- | --- |
| Intercept (I) | -15.68 | -13.1 | 9.66 | -16.54 |
| Body mass scaling species $i$ ($b$) | -0.25 | 0.25 | -0.45 | -0.31 |
| Body mass scaling predator ($c$) | | -0.8 | 0.47 | |
| Activation energy ($K$) | -0.84 | -0.38 | 0.26 | -0.69 |

### Food web structure, community properties and ecological stability

To assess the synergistic effects of temperature and invasions on food webs, we measured a set of statistical food web properties (Table 2), before and after invasions, across a range of temperatures. In addition to changes in food web properties we also assessed community properties such as total community biomass, the average species body size in the community (*AvgBS*), and the average ratio of predator vs. prey body masses (*AvgPPMR*). Lastly, to assess ecological stability, we focused on temporal variability of biomass both at the community and population levels. We quantified stability using two measures: (1) community invariability, measured as the inverse of total community biomass variability and (2) population invariability, as the inverse of average population-level biomass variability (Haegeman et al. 2016). Variabilities, both at the community (i.e. summing across the biomass of all species populations) and at the population level, were calculated as the ratio of the standard deviation to the mean biomass across the last 100 years of the simulations.

**Table 2.**
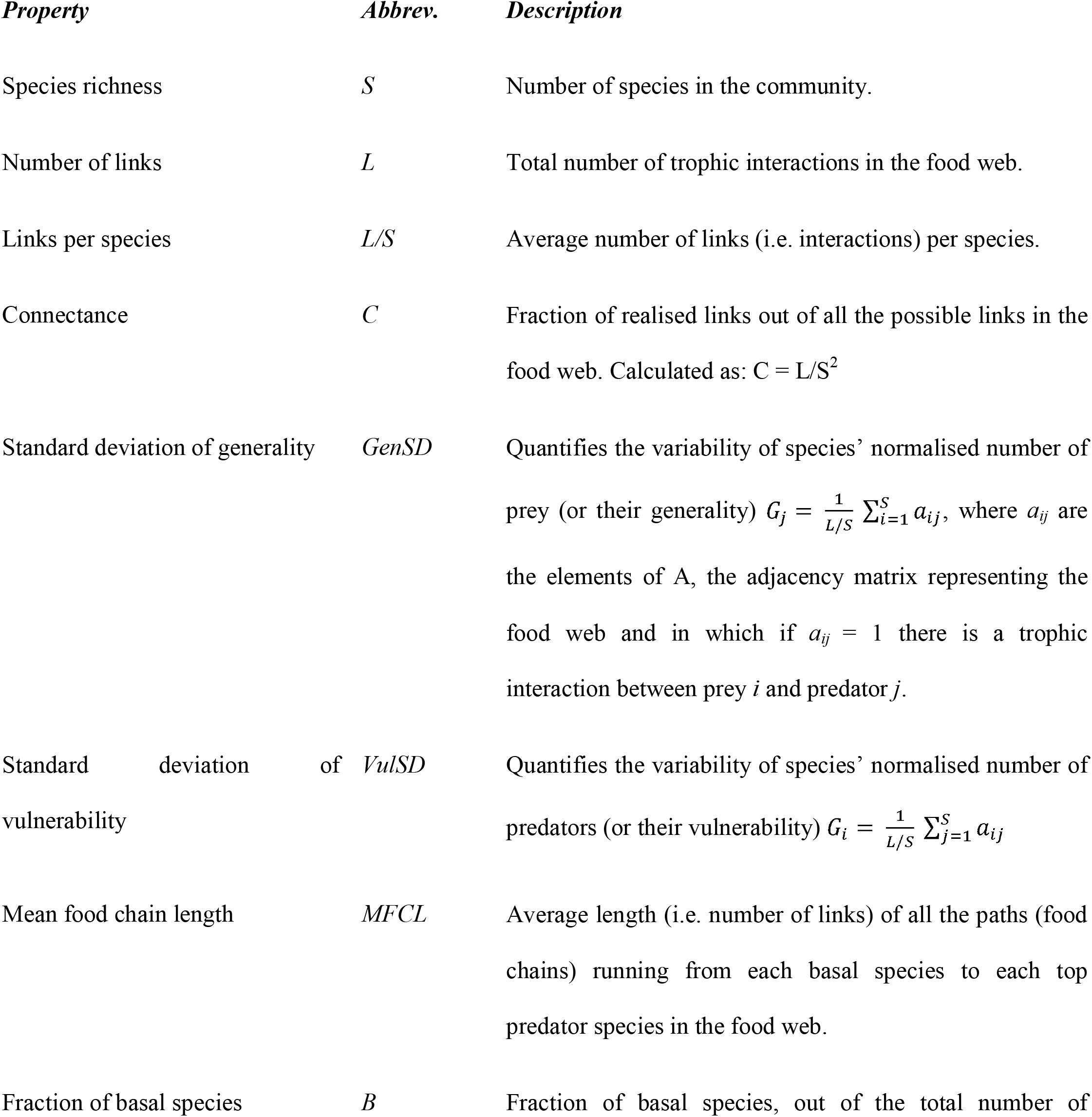
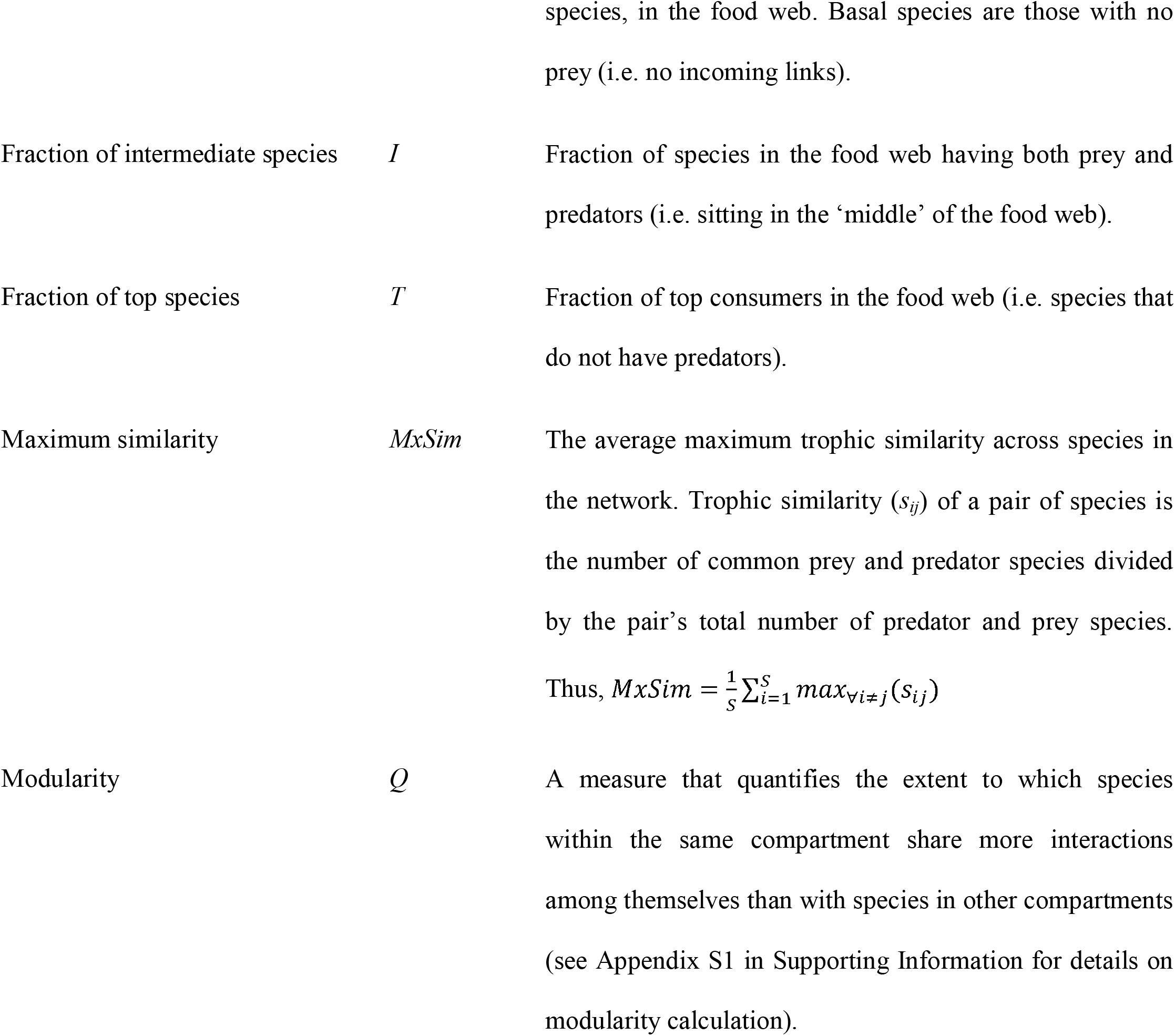
Network structural properties used in this study to quantify the structure of simulated food webs. The name of the properties, their abbreviations and a brief description are presented.

### Numerical simulations

Using the food web model specified above (Eqns. 1–5) we simulated a range of temperature regimes and the addition of new species (i.e. invasions) as follows:

1. 140 niche model food webs were randomly generated (*S* = 30 and *C* = 0.1).
2. Initial biomass densities for basal species were set to their carrying capacity *K* ≈ 2.75 following the allometric formulation in Eqn. 5b but omitting the temperature dependent term and assuming an allometric scaling constant and exponents of 10 and 0.28 respectively (Binzer *et al.* 2016). Consumer species initial abundances were set to 1/8 of this value as in Binzer *et al.* (2016).
3. Community dynamics were first simulated for an equivalent of 600 years (18.9216 × 10^9^ seconds) to achieve persistence (i.e. no further species extinctions) after initial transient dynamics. After these first 600 years, an equilibrium was reached and food web and community properties were calculated. The objective here was to quantify the direct effects of temperature on community structure before invasion. Resulting community features, and their relationship to temperature values, can then be related to invasion success.
4. After the first 600 years, an invasion was simulated by introducing a new species into the network. The introduced species was randomly drawn as an additional species from the original niche model network (i.e. ensuring it was different from all species originally present in the network). Since at this time point some species might have gone extinct (see step 6), rendering the potential introduced species disconnected, we repeated the drawing procedure if necessary until a connected species was found. This procedure avoided introducing an invasive species with no interactions.
5. After the introduction, we simulated further 600 years of network/community dynamics. We then recorded whether the invasion was successful (i.e. whether the introduced species persisted). Structural network and community properties were calculated again to assess community structure after the introduced species became invasive (i.e. established itself successfully in the community) or went extinct.
6. A species was considered extinct if at any point during the simulations its biomass fell below 10^−9^ g.m^−2^, at which point its abundance was set to 0.

For each of the 140 food webs, this procedure (i.e. steps 2 to 6 above) was repeated for each of 41 constant temperature regimes ranging from 0 to 40°C at 1°C intervals, yielding a total of 5740 numerical simulations. We used the same unique food webs for each temperature treatment to avoid confounding effects caused by initial differences in food web structure.

### Statistical analyses

The relationship between temperature and food web structure, stability and community properties - i.e. total biomass, *AvgBS*, and *AvgPPMR* -, and their corresponding effects on invasion success were analysed using piecewise structural equation models (SEMs). We computed SEMs considering invasion success (i.e. whether the invasive species established after introduction) as a response variable, with temperature affecting it directly and indirectly via network and community properties. The effect of species richness on food web and community properties was also incorporated into the SEMs, accounting thus for the indirect effect of temperature on network and community properties via *S* (see Appendix S2 for full SEM details).

To assess invasion effects on food web structure, stability and community properties, we calculated the ratio of values after (i.e. at the end of the simulations) vs. before (i.e. after initial transient dynamics) invasion for food web (Table 2) and community properties, as well as stability measures. To disentangle the direct and indirect effects (manifested via species richness) of temperature on these ratios, we performed another SEM following the same rationale as above and using the ratio of the effects as a response variable (i.e. effect size). We additionally assessed the differences between communities vulnerable vs. resistant to invasions in terms of these effects by comparing after/before ratios of each property between invaded and non-invaded communities using Mann-Whitney U tests.

All simulations and analyses were performed in R -language and environment for statistical computing (R Development Core Team 2017)-. Numerical simulations of ODEs were computed using the ode function of the deSolve package (Soetaert et al. 2010). Food web analyses were conducted with cheddar (Hudson et al. 2013). Modularity (Q) was computed using the cluster_louvain function from the igraph package (Csardi & Nepusz 2006). Piecewise SEMs and models within were performed using the piecewiseSEM (Lefcheck 2016) and the lme4 (Bates et al. 2015) packages respectively. Computer code developed to run model simulations and analyse outputs is available from the following repository: https://github.com/mlurgi/temperature-dependent-invasions.

## RESULTS

We focus on (i) the influence of warming on invasion success and (ii) the community-wide consequences of invasions. Effects of warming on food webs before invasion are detailed in Appendix S3. In line with previous findings (Binzer *et al.* 2016), we found that warmer communities harbour less species than their colder counterparts, particularly at high trophic levels, which in turn translates into higher connectance. These structural changes prompt an increase in both community biomass and stability in warmer environments (Appendix S3).

### How does warming influence invasion success?

Communities exposed to warmer temperatures were more prompt to invasions. Temperature had a direct positive effect on invasion success, although this effect was very weak (Fig. 1 and Table S1). This result holds even after accounting for temperature effects on species richness and other community and network properties (Table S1). Temperature influenced invasion success indirectly by modifying community properties, thus making communities more susceptible to invasions. Direct effects of network properties on invasion success were about an order of magnitude larger than the direct effect of temperature (compare arrow weights -i.e. standardised coefficients-on Fig. 1). In particular, the number of links (*L*), the average number of links per species (*L*/*S*), mean food chain length (*MFCL*) and the fraction of basal and intermediate species (*B* and *I* respectively) had a strong and significant influence on invasion success (Fig. 1 and Table S1). Surprisingly, we found no direct effect of species richness on invasion success (Table S1) which is likely due to the small variability in species richness in our communities after the initial transient dynamics (Fig. S1a and Appendix S3).

**Figure 1.**
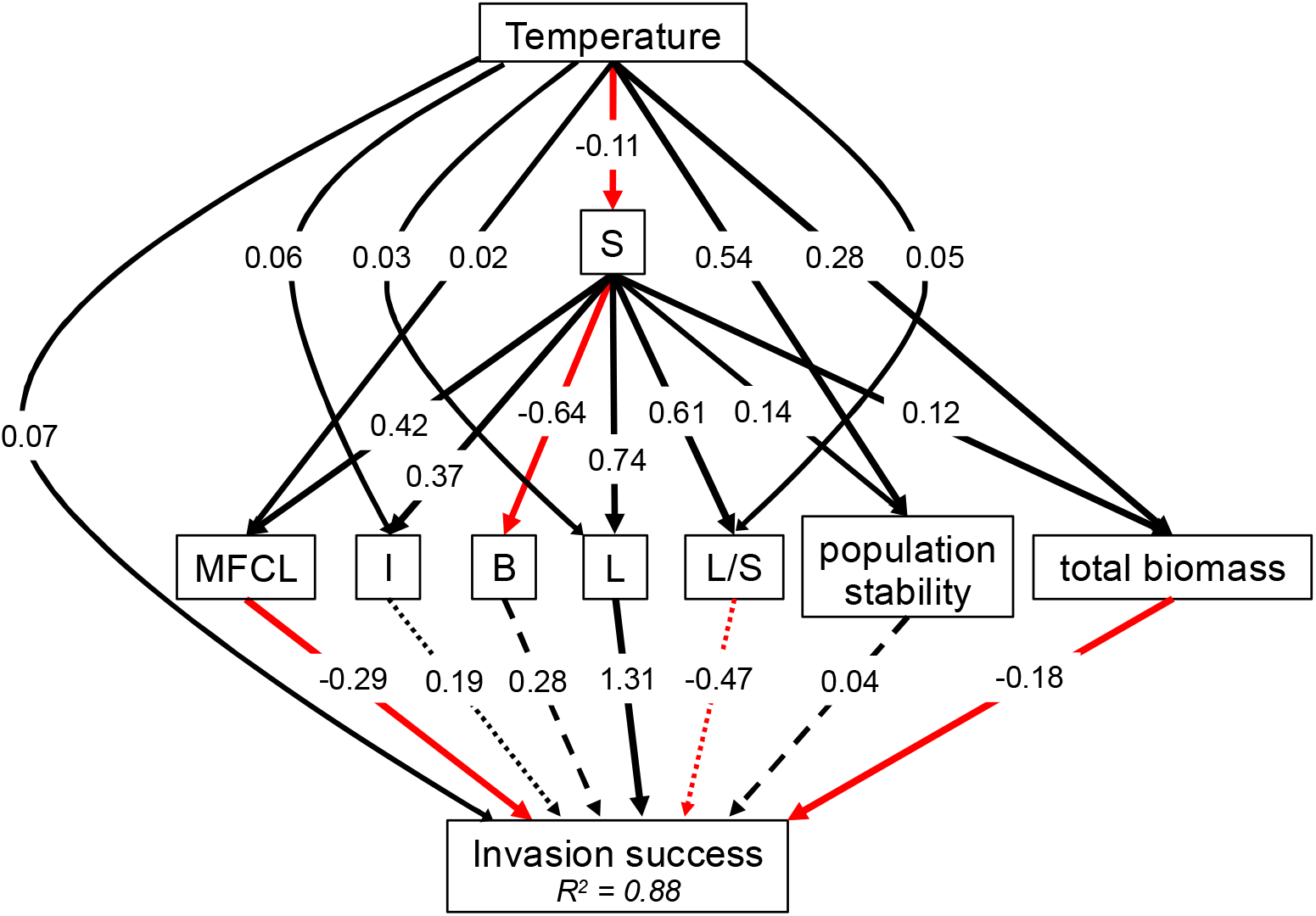
Structural equation model (SEM) describing the direct and indirect effects of temperature on invasion success in complex food webs. Arrows indicate the direct effects of predictor on response variables. Only predictors having a statistically significant effect (i.e. p-value < 0.05) on invasion success are shown (see Table S1 for more details). Black and red arrows represent positive and negative effects respectively. Solid, dashed, and dotted arrow styles represent strongly (p-value < 0.001), intermediate (p-value < 0.01), and marginally (p-value < 0.05) statistically significant effects respectively. Model fit: Fisher’s C = 22.67, p-value = 0.305, degrees of freedom = 20.

Even though warming had a direct significant effect on most network properties (Table S1, Figs. S1-S3), only a few of them affected invasion success. In particular, we found that communities with longer food chains (*MFCL*) were more resistant to invasion. In addition, communities with more links (*L*) and greater proportions of basal (*B*) and intermediate (*I*) species were more prompt to invasion (Fig. 1). Communities harbouring more specialised species (i.e. small *L*/*S*) also were more susceptible to invasion.

Changes in population stability and total community biomass also affected invasion success under warming. Whereas larger total community biomass conferred resistance against invasions, communities with higher population stability were easier to invade (Fig. 1). Overall our results show that indirect effects of temperature on invasion success, mediated by changes in network and community properties and dynamics, were stronger than direct ones.

### Ecological consequences of invasions along the temperature gradient

Overall, invasions strongly decreased species richness (Fig. 2a), which, in turn, affected several network properties. Moreover, we found that the magnitude of the change of food web and community properties driven by invasions often depended on temperature (Fig. 2 & Table S2). In particular, warmer communities loose more species and interactions when invaded than their colder counterparts. This translates into more connected communities (Fig. 2a). Higher connectance (*C*) was accompanied by a larger heterogeneity in the distribution of predators among prey species (V*ulSD*) and a stronger increase in the fraction of top predators (*T*). On the other hand, warming prompted a stronger decrease in the proportion of intermediate species (*I*) when invaded. The proportion of basal species (B) was not influenced by invasion or temperature (Table S2). This suggests that intermediate species became top predators when their predators disappeared in invaded communities, yielding a lower fraction of intermediates while increasing the fraction of top predator species. Warming also lead to a stronger decrease in modularity (*Q*) which is explained by the presence of a larger fraction of top consumers and a higher connectance.

**Figure 2.**
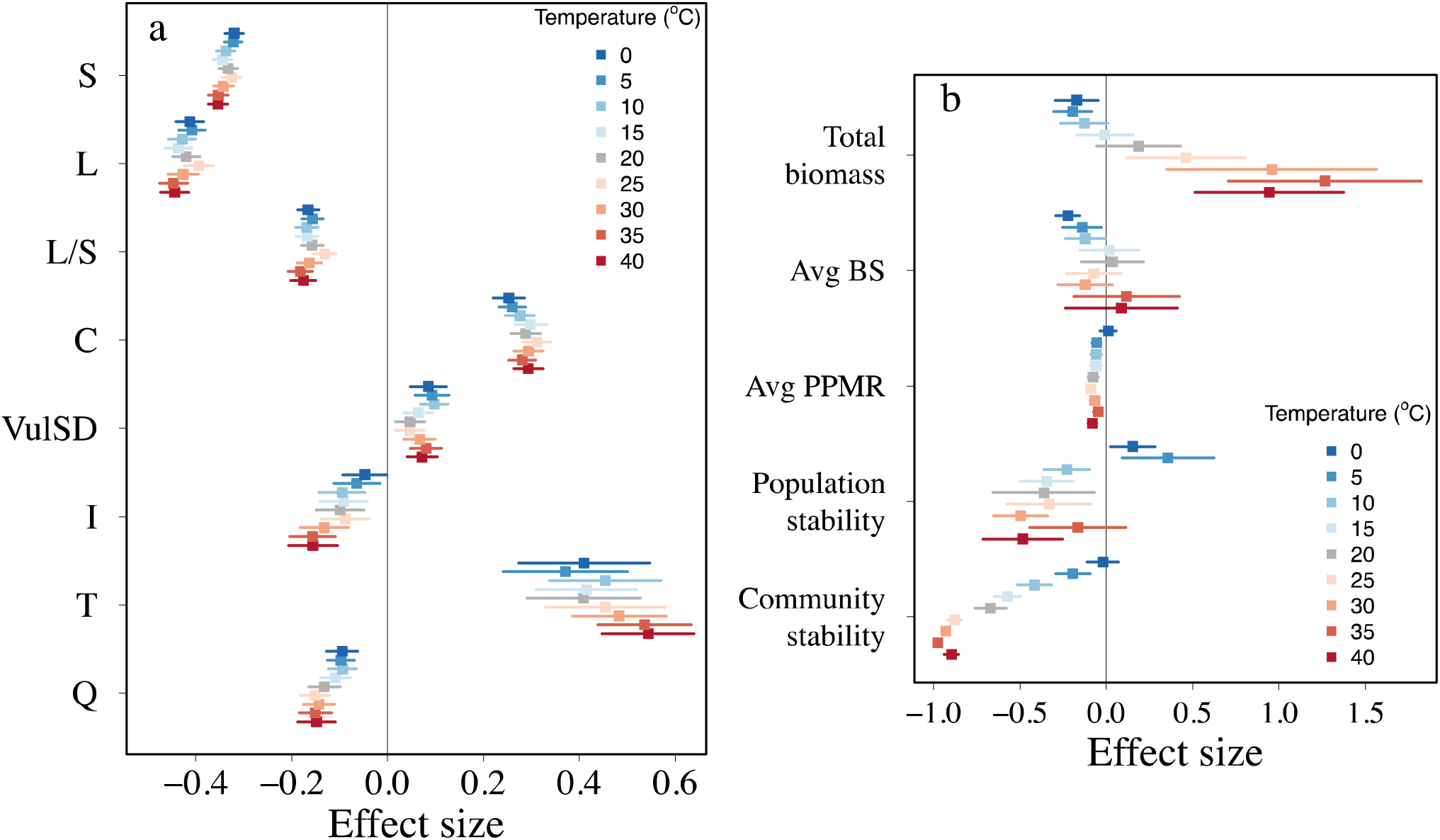
Effect sizes (mean ± s.d.) of successful invasions on complex food webs across temperature regimes on network (a) and community (b) properties. Effect sizes were quantified as the ratio between the values of the network/community property after species introduction vs. before introduction, minus unity. Negative values thus indicate negative effects of the invasion on the community (i.e. the value after the invader’s establishment is smaller than before the introduction). Only effects on properties identified by SEMs as being significantly influenced by temperature (Table S2) are shown. Only a subset of temperature regimes is shown to ease the visualisation of the results.

Community properties and stability were affected more heterogeneously by invasions across temperature regimes than network structure (Fig. 2). Total community biomass increased in warm invaded communities relative to their pre-invasion state (positive effect sizes), whereas it decreased under colder conditions (Fig. 2b, negative effect sizes). Similarly, population stability increased in colder communities (at 0 and 5°C) but decreased in warmer ones (although the effect was weak at 35°C). In addition, invasions always decreased community stability (Fig. 2b), although this effect was weaker in colder environments compared to warmer ones. Overall, we found that warm invaded communities have less species, with these species fluctuating more over time (i.e. reduced stability) than cold invaded communities.

Lastly, the average body size (*AvgBS*) of species in the community remains mostly unaffected by invasions except at very low temperatures where invasions decreased average body size. On the other hand, the average predator:prey body size ratio (*AvgPPMR*) was negatively impacted by invasion and this effect was more pronounced when temperature increased (Fig. 2b). The decrease in body size ratio, along with the considerable increase in the fraction of top predator species (Fig. 2a), while the fraction of basal species was unaffected by invasion or warming reinforces the observation that top predators were lost and replaced by consumers further down the food web. Interestingly, this replacement appeared to be stronger in warmer communities that also lost more species than colder ones.

The effects of unsuccessful invasions (i.e. when the introduced species went extinct) on food webs were more homogeneous across the temperature gradient than those caused by successful invasions, mainly affecting species numbers, connectivity (*L* and *L/S*), and the fraction of intermediate species (Appendix S4).

## DISCUSSION

Global warming and biological invasions affect communities simultaneously. It is fundamental to better understand their synergistic effects on biodiversity (Bradley *et al.* 2010). Using dynamical models we have investigated the interactive effects of warming and invasions on the structure and dynamics of complex food webs. We showed that warming has two overall effects on invasions and on invaded communities. First, it modified key aspects of community structure and dynamics that in turn facilitated invasions by introduced species. Secondly, warming mostly amplified the impacts of invasions on the structure and organisation of communities. Importantly, the directionality of the effects of invasions on recipient communities changed across the temperature gradient for community biomass and population stability. Further, our results suggest that the direct effects of temperature on invasion success are weaker to those mediated by changes in community structure and stability.

Previous attempts to understand and predict the combined effects of warming and invasions have been based mainly on bioclimatic envelop models, relying on temperature thresholds for survival and reproduction of the invasive species (Stachowicz *et al.* 2002; Walther *et al.* 2009; Sorte *et al.* 2010; Kent *et al.* 2018). Such phenomenological approaches, even though informative, lack a mechanistic understanding of how warming mediates invasions and how both can synergistically affect ecological communities. Here, we have provided a first step towards a better understanding of the synergistic effects of warming and invasions on complex ecological communities. Our efforts complement recent attempts to better understand how warming modulates the effects of invasions on natural communities (Zhang et al. 2017).

### Warming effects on invasion success

We found that, before invasion, warming increased food web stability but decreased the persistence of top predators, as reported in previous studies (Fussmann *et al.* 2014; Binzer *et al.* 2016; Sentis *et al.* 2017). This increased stability is likely explained by average trophic interaction strength decreasing and by consumers being less efficient at feeding relative to their metabolic losses (Boukal *et al.* 2019). Warmer communities thus contained less species, but network connectance increased. Whether these warming-induced changes influence invasion success remains an open question.

Previous theoretical models showed that more connected food webs are generally better at repelling invaders (Romanuk *et al.* 2009). In contrast, Lurgi *et al.* (2014) showed that, when controlling for the number of species, less connected food webs are more resistant to invasions. Here, we found that invasions are more successful in warmer communities, which are more connected but poorer in species richness than colder ones. Invasion success was strongly mediated by the indirect effects of temperature on network and community properties and stability, with these effects only weakly mediated by species richness. Temperature had an impact on the composition of species across trophic levels, influencing greatly the proportion of top predators. This, in turn, made communities more susceptible to invasions by reducing mean food chain length. We also found that warming directly increased invasion success. Nevertheless, the direct effect of warming on invasion success was much weaker than its indirect effects.

Our results are in line with empirical studies reporting that the influence of warming on invasion success may depend on how warming affects the strength of trophic interactions and the persistence of local predators or competitors (Holzapfel & Vinebrooke 2005; Lu *et al.* 2013; Seifert *et al.* 2015). Warming can prevent invasions by increasing top-down control on introduced species (Lu *et al.* 2013; Lu *et al.* 2016). On the other hand, Holzapfel and Vinebrooke (2005) showed that warming can enhance invasions by removing top-down control on the invader following predator extinctions. Our model results suggest that the loss of top predators, which in turn reduces mean food chain length, is a driving mechanism of invasion success. This highlights the importance of species loss at high trophic levels on facilitating invasions under warming.

### Ecological consequences of invasions along a temperature gradient

Our results are in line with previous studies showing that invasions decrease species richness and, as a consequence, impact strongly community structure (Hui & Richardson 2019). However, recent theoretical studies have suggested that warming is predicted to have an opposite effect on invasions, by enhancing species richness and ecosystem functioning (Zhang et al. 2017). We found that temperature can modulate the consequences of invasions for community structure. Warmer communities tend to lose more species and interactions when invaded, which translates into more connected communities compared to colder ones. This intensification in the effects of invasions across the temperature range highlights the importance of considering a full spectrum of temperature treatments to fully understand the effects of warming on invaded communities.

The proportion of top predator species in our model food webs increased as a consequence of invasion, especially in warmer environments. This result apparently contradicts the loss of top predators following invasions that we observed. Both observations are reconciled by a third result: invasions shorten food chains. In short, when top predators go extinct they are replaced by consumers further down the food web, which in turn become top predators. This switch decreases the proportion of intermediate species while increasing the proportion of top species, ultimately shortening food chain length. Invasions thus exacerbate the previously observed effect of warming on top predator species, and corroborates previous empirical findings of higher trophic levels being most vulnerable to climate change (Voigt *et al.* 2003). We should thus expect warmer and invaded communities being even more susceptible to invasions, entering a positive feedback loop via the loss of predator species.

As any modelling study, ours relies on a set of assumptions that can influence model predictions. For instance, we did not account for temperature fluctuations, evolutionary change or differences in the thermal traits of the invasive species (e.g. the invasive species may have thermal traits better adapted to warmer climate) that can also influence invasion success and species persistence (Vasseur *et al.* 2014; Zhang *et al.* 2017). The assumption of invasive species possessing similar thermal performance than resident species is supported by the observation that most invasive species are introduced by human transport and their invasion success is not proven to be strongly related to the climate in their native environment (Gippet *et al.* 2019). Additionally, in our model, introduced species, and their biotic interactions, have been defined following the same heuristics as the rest of the species in the recipient community (i.e. they are drawn from the niche model), whereas in natural communities invasive species tend to be generalists, both in terms of habitats and interactions (Elton 1958). We decided to define introduced species in this way to be as parsimonious as possible in our experiments and avoid the confounding effects of over-generalist species in modelled communities. It would be interesting to extend the results presented here using invasive species with more realistic traits. Despite these limitations, our model provides a first step in the exploration of the consequences of warming and invasions in species-rich communities.

## CONCLUSION

Research on the effects of temperature on biological invasions has traditionally focused on the species level and have not explicitly considered species interactions, usually not taking into account the way temperature can affect food web and community properties prior to invasions. Understanding the joint effects of warming and invasions at the community level constitutes a pressing challenge to unveil the full consequences of global change on natural communities. Here we addressed this challenge and showed that temperature’s direct effects on invasion success are weaker than its indirect effects mediated by changes in food web structure, community properties and stability. Moreover, we showed that the impact of invasions depend on the temperature experienced by the invaded communities. Warmer food webs lose more species and interactions when invaded than their colder counterparts. These changes are accompanied by an increase in the fraction of top predator species, enhanced total community biomass and decreased stability. Overall, our study suggest that both warming and invasion act synergistically to fasten species loss creating smaller and more connected networks. It paves way for a better understanding of the causes and consequences of invasions in a warming world.

## ACKNOWLEDGEMENTS

This work was funded by the FRAGCLIM Consolidator Grant (number 726176) from the European Research Council under the European Union’s Horizon 2020 Research and Innovation Program. ML was additionally supported by the French ANR through LabEx TULIP (ANR-10-LABX-41; ANR-11-IDEX-002-02), and by a Region Midi-Pyrénées Project (CNRS 121090) to JMM. AS was additionally supported by the ANR project EcoTeBo (ANR-19-CE02-0001-01) from the French National Research Agency (ANR).

