## Appendix for "Warming indirectly increases invasion success in food webs" (from another version of this preprint: version 1, 2020-07-21)

**Supporting Information**

**Appendix S1. *Supplementary Methods: Modularity calculation***

Modularity is a measure used to quantify the extent to which networks are organised into modules (i.e. compartments) within which links (i.e. interactions) amongst species are significantly more frequent than with species sitting outside (i.e. in other modules), within the same network. To identify modules within the network, and hence be able to quantify modularity, an optimisation algorithm to obtain optimal partitions of nodes into modules, aka community detection algorithms, must be used. We used the Louvain optimisation algorithm for community detection (Blondel *et al.* 2008). This algorithm uses *Q* (Eq. 1) as a measure of modularity (Girvan & Newman 2002).

$Q=\frac{1}{2m}\sum_{i,j} \left( A_{ij}-\frac{k_{i}k_{j}}{2m} \right)\delta\left( g_{i},g_{j} \right)$	(1)

where *m* is the number of links in the network, *A_ij_* are the elements of the adjacency matrix representing the network of interactions (1 if there is a link between species *i* and *j*, and 0 otherwise), *k_i_* and *k_j_* are the total number of links of species *i* and *j* respectively, *g_i_* is the module to which species *i* belongs; and $\delta$ is a function such that $\delta\left( r,s \right)$ = 1 if *r* = *s* and 0 otherwise.

**Appendix S2. *Supplementary Methods: Building structural equation models***

Structural Equation Models (SEMs) are a type of path analysis in which the effects of a predictor on a response variable can be modelled directly and indirectly through the action of additional variables. Using SEMs, we can thus disentangle the direct and indirect effects of temperature on invasion success mediated by changes in species biological rates (direct thermal effects) or in community properties (indirect thermal effects). SEMs require the modelling of each relationship (i.e. path) as a regression model and are thus composed of multiple regression models. All variables considered (except invasion success, which takes values of 0 and 1) were standardised (i.e. 0 mean and 1 standard deviation) before proceeding with the SEM modelling and analysis. This facilitates comparison of estimated coefficients of the relationships across predictor variables and across different paths in the SEM. To define the SEM predicting invasion success from temperature and community properties (i.e. food web and community properties and stability measures, see *Material and Methods* in main text for all properties considered), we used linear mixed effects (LMEs) models for the relationships between temperature and each of these properties. We also included a LME for the relationship between species richness and each community property because of the known effect of species richness on food web and other community properties (Dunne *et al.* 2002). An identifier for each of the 140 generated networks (ID) was included as a random effect to account for the replication of each network across temperature regimes. To investigate the effects of the community properties and temperature on invasion success (i.e. a discrete binomial variable), we used generalised LMEs (GLMEs) with network ID as random effect and with binomial distribution family. LMEs and GLMEs included in the SEMs were thus defined as to establish all potential paths from temperature through food web / community structure and stability to invasion success. All independence claims, between response variables of the different models, identified as significant by an initial fit of the SEMs were then added to the SEM specification to specify that these variables are allowed / expected to freely covary (Y1 %~~% Y2).

To disentangle the direct influence of temperature on the effects of invasions from those indirect effects brought about by changes on species richness and number of links, another SEM was constructed. In this case responses variables were defined as the effect of invasion on each network and community property (i.e. ratio before vs. after invasion, see *Material and Methods* in main text). Predictor variables included were thus: temperature, number of species before invasion (*Original S*), and the effects of the invasion on both *S* and *L*.

**Appendix S3*. Supplementary Results: Warming effects on food webs before invasion***

We found that warming decreased species richness while connectance displayed the opposite pattern (Fig. S1b). In both cases, these effects were relatively weak as, on average, only two species went extinct with warming and connectance increased by only 0.01 over the thermal gradient. We also found opposite patterns for mean food chain length and generality with the former mostly decreasing with warming and the later mostly increasing with warming. Finally, both modularity and vulnerability decreased with warming. Interestingly, for all of these 6 structural properties (except vulnerability) the effect of temperature was stronger and linear for temperatures below 20°C whereas it was weaker for temperatures above 20°C.

This indicates that warming has non-linear effects on community structure and that warming has the strongest effect for communities inhabiting cooler environments. We found that warming influences the proportion of basal, intermediate and top species within the food webs (Fig. S2): it decreased the proportion of top species while increasing basal species proportion. Interestingly, the proportion of intermediate species remained fairly constant across the temperature range. Overall, warmer communities are smaller (i.e. have less species), more connected, and ‘shorter’ (i.e. shorter food chains). They achieve this by maintaining a constant number of intermediate species while increasing the proportion of basal at the expense of top species.

The decrease in vulnerability as temperature increases, accompanied by the increase in generality in the light of a smaller fraction of top species and larger connectivity suggests that prey/resource species are being consumed by more species in general while more generalist consumers appear in warmer communities. We found that both community and population level stabilities as well as connectivity and biomass increase with warming. This suggests that, by loosing top predators and increasing connectivity, warmer food webs increase in biomass, powered by enhanced productivity and consumption rates.

These observations are in line with previous studies showing that warming increases food web stability but decreases the persistence of top species (Fussmann *et al.* 2014; Binzer *et al.* 2016; Sentis *et al.* 2017). The increased stability is likely explained by average trophic interaction strength decreasing due to consumers being less efficient at feeding relative to their metabolic losses (Boukal *et al.* 2019). As a consequence, warmer communities are composed of less species but these species are more connected, with the resulting communities being more stable.

**Appendix S4*. Supplementary Results: Are unsuccessful introductions less harmful?***

We found that the effects of unsuccessful introductions across temperature regimes affected almost exclusively network structure, with community properties being affected in a similar way to what was observed for successful invaders (Table S3). The effects of unsuccessful introductions on the proportion of top predator species (*T*), the vulnerability (*Vul SD*), and modularity (*Q*), were indistinguishable from those brought by successful invasions. For the remaining properties, even though the effects were in the same general directions (Fig. S4), they were stronger for unsuccessful invasions than for successful ones (compare Figs. 2a and S4). These stronger effects might be the very reason why the invasions were unsuccessful. These results suggest that regardless of whether introduced species manage to establish themselves in recipient communities they can have profound impacts on community organisation. These effects are comparable to, and in some cases even stronger than, those brought about by successful invaders.

**Table S1*. Direct and indirect effects of temperature on invasion success.*** The outcome of piecewise structural equation models (SEMs) show the significance (p.value) and strength (estimate) of the relationship that each predictor variable has on each response variable. See methods for a full description of the abbreviated names of network properties. Entries in the table are ordered by the magnitude of the estimate of each predictor-response pair.

| ***Response*** | ***Predictor*** | ***Std. Estimate*** | ***Std. Error*** | ***p value*** |
| --- | --- | --- | --- | --- |
| Invasion success | Temperature | 0.0695 | 0.0858 | < 0.001 |
| Invasion success | *S* | -0.2074 | 0.7143 | 0.1187 |
| Invasion success | *L* | 1.3066 | 1.3758 | < 0.001 |
| Invasion success | *L/S* | -0.4736 | 1.0474 | 0.0151 |
| Invasion success | *MFCL* | -0.2856 | 0.3468 | < 0.001 |
| Invasion success | *Q* | 0.058 | 0.1928 | 0.106 |
| Invasion success | *GenSD* | -0.0844 | 0.4362 | 0.2985 |
| Invasion success | *VulSD* | 0.0762 | 0.23 | 0.0751 |
| Invasion success | community stab. | 0.0127 | 0.0581 | 0.2395 |
| Invasion success | population stab. | 0.0345 | 0.0687 | 0.0069 |
| Invasion success | *Avg PPMR* | -0.0496 | 0.2022 | 0.1876 |
| Invasion success | Total biomass | -0.1842 | 0.1545 | < 0.001 |
| Invasion success | *B* | 0.2747 | 0.5115 | 0.0039 |
| Invasion success | *I* | 0.1896 | 0.4488 | 0.0232 |
| Invasion success | *MaxSim* | 0.0186 | 0.4283 | 0.8159 |
| *S* | Temperature | -0.1141 | 0.0038 | < 0.001 |
| *L* | *S* | 0.7432 | 0.0042 | < 0.001 |
| *L* | Temperature | 0.0289 | 0.0013 | < 0.001 |
| *L/S* | *S* | 0.6138 | 0.0077 | < 0.001 |
| *L/S* | Temperature | 0.0465 | 0.0024 | < 0.001 |
| *MFCL* | *S* | 0.4211 | 0.0084 | < 0.001 |
| *MFCL* | Temperature | 0.0244 | 0.0026 | < 0.001 |
| *Q* | *S* | 0.0096 | 0.0148 | 0.5185 |
| *Q* | Temperature | -0.0412 | 0.0046 | < 0.001 |
| *GenSD* | *S* | -0.418 | 0.0104 | < 0.001 |
| *GenSD* | Temperature | 0.0183 | 0.0032 | < 0.001 |
| *VulSD* | *S* | 0.4797 | 0.0168 | < 0.001 |
| *VulSD* | Temperature | -0.0767 | 0.0052 | < 0.001 |
| community stab. | *S* | -0.0924 | 0.0284 | 0.0013 |
| community stab. | Temperature | -0.2591 | 0.0121 | < 0.001 |
| population stab. | *S* | 0.1404 | 0.0234 | < 0.001 |
| population stab. | Temperature | 0.5416 | 0.0105 | < 0.001 |
| *Avg PPMR* | *S* | -0.7147 | 0.0149 | < 0.001 |
| *Avg PPMR* | Temperature | -0.0544 | 0.0047 | < 0.001 |
| Total biomass | *S* | 0.1247 | 0.0158 | < 0.001 |
| Total biomass | Temperature | 0.2764 | 0.005 | < 0.001 |
| *B* | *S* | -0.6365 | 0.0072 | < 0.001 |
| *B* | Temperature | 0.0006 | 0.0022 | 0.7808 |
| *I* | *S* | 0.3768 | 0.0083 | < 0.001 |
| *I* | Temperature | 0.0574 | 0.0026 | < 0.001 |
| *MaxSim* | *S* | 0.4411 | 0.0091 | < 0.001 |
| *MaxSim* | Temperature | 0.0653 | 0.0028 | < 0.001 |

**Table S2*. Direct and indirect effects of temperature on changes in network and community properties after successful invasions.*** The outcome of piecewise structural equation models (SEMs) show the significance (p.value) and strength (estimate) of the relationship that each predictor variable has on each response variable. See methods for a full description of the abbreviated names of network properties. In the table, network and community properties refer to the effect of invasions on that variable (measured as the ratio between the value of that property after vs. before invasion), except for *Original S* which refer to the actual values of number of species before invasion. All variables were standardised (0 mean and 1 SD) before analysis to make estimated coefficients comparable across relationships. Entries in the table are ordered by the magnitude of the estimate of each predictor-response pair.

| ***response*** | ***predictor*** | ***estimate*** | ***std.error*** | ***p.value*** |
| --- | --- | --- | --- | --- |
| *C* | *S* | -2.5003024 | 0.02653286 | <0.001 |
| *C* | *L* | 2.25452757 | 0.02568152 | <0.001 |
| *L/S* | *L* | 1.72506638 | 0.01318993 | <0.001 |
| *Q* | *S* | 1.45480538 | 0.04332444 | <0.001 |
| *Q* | *L* | -1.4259227 | 0.04191024 | <0.001 |
| *MaxSim* | *L* | 1.07029126 | 0.02753079 | <0.001 |
| *L* | *S* | 0.9424 | 0.0083 | <0.001 |
| *L/S* | *S* | -0.8422227 | 0.01365003 | <0.001 |
| *MFCL* | *L* | 0.68066601 | 0.0305 | <0.001 |
| *I* | *L* | 0.66025887 | 0.03772995 | <0.001 |
| *B* | *L* | -0.5835298 | 0.03357543 | <0.001 |
| *Total biomass* | *L* | -0.506669 | 0.06927665 | <0.001 |
| *MaxSim* | *S* | -0.4978979 | 0.02851559 | <0.001 |
| *Avg PPMR* | *L* | -0.472161 | 0.04434102 | <0.001 |
| *population stability* | *S* | -0.3848517 | 0.08003016 | <0.001 |
| *GenSD* | *L* | -0.3620868 | 0.03135835 | <0.001 |
| *I* | *Original S* | 0.32881045 | 0.03160832 | <0.001 |
| *community stability* | *Temperature* | -0.3268959 | 0.01576983 | <0.001 |
| *Total biomass* | *S* | 0.31739092 | 0.07159196 | <0.001 |
| *T* | *L* | -0.313287 | 0.0404939 | <0.001 |
| *community stability* | *Original S* | 0.28820071 | 0.04824158 | <0.001 |
| *VulSD* | *Original S* | -0.2809838 | 0.04176833 | <0.001 |
| *population stability* | *Original S* | 0.27466547 | 0.05734222 | <0.001 |
| *I* | *S* | -0.2649172 | 0.03902797 | <0.001 |
| *Avg BS* | *S* | 0.24851865 | 0.02747021 | <0.001 |
| *VulSD* | *L* | -0.2303383 | 0.05199867 | <0.001 |
| *Avg PPMR* | *S* | 0.18767334 | 0.04589615 | <0.001 |
| *GenSD* | *S* | 0.18104094 | 0.03247663 | <0.001 |
| *S* | *Original S* | -0.1675601 | 0.03831307 | <0.001 |
| *Total biomass* | *Temperature* | 0.15472442 | 0.01638982 | <0.001 |
| *MFCL* | *Original S* | -0.1500543 | 0.02579755 | <0.001 |
| *C* | *Original S* | 0.14557254 | 0.02048923 | <0.001 |
| *MaxSim* | *Original S* | 0.13658943 | 0.02368916 | <0.001 |
| *population stability* | *Temperature* | -0.1236577 | 0.01850394 | <0.001 |
| *Original S* | *Temperature* | -0.0954658 | 0.00494403 | <0.001 |
| *VulSD* | *Temperature* | -0.089599 | 0.01230228 | <0.001 |
| *S* | *Temperature* | -0.053705 | 0.01084818 | <0.001 |
| *L/S* | *Original S* | 0.05168671 | 0.01107052 | <0.001 |
| *I* | *Temperature* | -0.0420687 | 0.00894355 | <0.001 |
| *L/S* | *Temperature* | 0.02224161 | 0.00312086 | <0.001 |
| *Avg BS* | *Original S* | -0.1873257 | 0.04635507 | <0.001 |
| *Avg PPMR* | *Original S* | 0.14314037 | 0.03741667 | <0.001 |
| *C* | *Temperature* | 0.02362991 | 0.00607648 | <0.001 |
| *T* | *Original S* | -0.1232746 | 0.03354702 | <0.001 |
| *GenSD* | *Original S* | -0.0904746 | 0.02690122 | <0.001 |
| *Q* | *Original S* | 0.1084267 | 0.03403441 | 0.0015 |
| *Avg PPMR* | *Temperature* | 0.03335829 | 0.0104924 | 0.0015 |
| *Total biomass* | *Original S* | -0.1706438 | 0.05572255 | 0.0023 |
| *population stability* | *L* | 0.22486487 | 0.07759341 | 0.0038 |
| *community stability* | *S* | -0.1866029 | 0.06438162 | 0.0038 |
| *L* | *Temperature* | -0.0322267 | 0.0112336 | 0.0042 |
| *T* | *Temperature* | 0.02680508 | 0.00957986 | 0.0052 |
| *B* | *S* | 0.08942478 | 0.03475715 | 0.0102 |
| *Avg BS* | *Temperature* | 0.03974613 | 0.01565981 | 0.0113 |
| *T* | *S* | -0.104488 | 0.04188799 | 0.0127 |
| *Q* | *Temperature* | -0.0233157 | 0.00991465 | 0.0188 |
| *community stability* | *L* | 0.14382574 | 0.06236997 | 0.0213 |
| *MFCL* | *Temperature* | -0.0123556 | 0.00721751 | 0.0871 |
| *B* | *Original S* | 0.04850508 | 0.02843158 | 0.0889 |
| *L* | *Original S* | -0.0209 | 0.0156 | 0.1819 |
| *B* | *Temperature* | 0.01226166 | 0.00794546 | 0.123 |
| *MFCL* | *S* | -0.0252228 | 0.03157223 | 0.4246 |
| *MaxSim* | *Temperature* | 0.00428173 | 0.00651754 | 0.5114 |
| *VulSD* | *S* | 0.03265217 | 0.05373417 | 0.5438 |
| *GenSD* | *Temperature* | 0.00370016 | 0.00742304 | 0.6183 |

**Table S3*. Difference in the effects on network/community structure of successful vs. unsuccessful invasions.*** Results of Mann Whitney U tests comparing the effects on species introductions, measured as the ratio of community/network properties after transient dynamics following introduction vs. before introduction across temperature treatments, shown here at 5 °C intervals for clarity. Comparisons were made for all properties.

| ***Network / community property*** | ***Temp. (ºC)*** | ***Median invaded*** | ***Median not invaded*** | ***U*** | ***Effect size*** | ***n_1_*** | ***n_2_*** | ***p value*** |
| --- | --- | --- | --- | --- | --- | --- | --- | --- |
| *S* | 0 | 0.68 | 0.6 | 3051.5 | 0.62 | 67 | 73 | 0.012 |
| *S* | 5 | 0.67 | 0.58 | 3274 | 0.67 | 68 | 72 | 0.001 |
| *S* | 10 | 0.66 | 0.56 | 3327 | 0.68 | 66 | 74 | <0.001 |
| *S* | 15 | 0.65 | 0.57 | 3080 | 0.63 | 63 | 77 | 0.006 |
| *S* | 20 | 0.68 | 0.57 | 3263 | 0.68 | 61 | 79 | <0.001 |
| *S* | 25 | 0.68 | 0.55 | 3375 | 0.7 | 60 | 80 | <0.001 |
| *S* | 30 | 0.68 | 0.55 | 3207.5 | 0.67 | 61 | 79 | 0.001 |
| *S* | 35 | 0.65 | 0.56 | 3158 | 0.65 | 65 | 75 | 0.003 |
| *S* | 40 | 0.65 | 0.55 | 3220.5 | 0.67 | 61 | 79 | 0.001 |
| *L* | 0 | 0.57 | 0.5 | 3115.5 | 0.64 | 67 | 73 | 0.005 |
| *L* | 5 | 0.57 | 0.45 | 3311.5 | 0.68 | 68 | 72 | <0.001 |
| *L* | 10 | 0.57 | 0.4 | 3400.5 | 0.7 | 66 | 74 | <0.001 |
| *L* | 15 | 0.57 | 0.42 | 3101 | 0.64 | 63 | 77 | 0.005 |
| *L* | 20 | 0.59 | 0.42 | 3291 | 0.68 | 61 | 79 | <0.001 |
| *L* | 25 | 0.62 | 0.39 | 3539.5 | 0.74 | 60 | 80 | <0.001 |
| *L* | 30 | 0.55 | 0.4 | 3285.5 | 0.68 | 61 | 79 | <0.001 |
| *L* | 35 | 0.55 | 0.42 | 3234 | 0.66 | 65 | 75 | 0.001 |
| *L* | 40 | 0.53 | 0.4 | 3319 | 0.69 | 61 | 79 | <0.001 |
| *L/S* | 0 | 0.88 | 0.81 | 3106 | 0.64 | 67 | 73 | 0.006 |
| *L/S* | 5 | 0.86 | 0.79 | 3245 | 0.66 | 68 | 72 | 0.001 |
| *L/S* | 10 | 0.86 | 0.72 | 3354 | 0.69 | 66 | 74 | <0.001 |
| *L/S* | 15 | 0.86 | 0.78 | 3017 | 0.62 | 63 | 77 | 0.013 |
| *L/S* | 20 | 0.87 | 0.75 | 3137 | 0.65 | 61 | 79 | 0.002 |
| *L/S* | 25 | 0.89 | 0.7 | 3492.5 | 0.73 | 60 | 80 | <0.001 |
| *L/S* | 30 | 0.88 | 0.71 | 3208 | 0.67 | 61 | 79 | 0.001 |
| *L/S* | 35 | 0.86 | 0.72 | 3127 | 0.64 | 65 | 75 | 0.004 |
| *L/S* | 40 | 0.89 | 0.73 | 3293 | 0.68 | 61 | 79 | <0.001 |
| *C* | 0 | 1.21 | 1.2 | 2605 | 0.53 | 67 | 73 | 0.507 |
| *C* | 5 | 1.26 | 1.27 | 2347 | 0.48 | 68 | 72 | 0.675 |
| *C* | 10 | 1.29 | 1.26 | 2415 | 0.49 | 66 | 74 | 0.912 |
| *C* | 15 | 1.31 | 1.32 | 2313 | 0.48 | 63 | 77 | 0.639 |
| *C* | 20 | 1.31 | 1.29 | 2308.5 | 0.48 | 61 | 79 | 0.673 |
| *C* | 25 | 1.3 | 1.24 | 2492.5 | 0.52 | 60 | 80 | 0.698 |
| *C* | 30 | 1.29 | 1.24 | 2345 | 0.49 | 61 | 79 | 0.788 |
| *C* | 35 | 1.28 | 1.26 | 2232 | 0.46 | 65 | 75 | 0.392 |
| *C* | 40 | 1.26 | 1.25 | 2423.5 | 0.5 | 61 | 79 | 0.955 |
| *Avg BS* | 0 | 0.82 | 0.67 | 2663 | 0.54 | 67 | 73 | 0.365 |
| *Avg BS* | 5 | 0.81 | 0.27 | 3157 | 0.64 | 68 | 72 | 0.003 |
| *Avg BS* | 10 | 0.92 | 0.38 | 2994 | 0.61 | 66 | 74 | 0.021 |
| *Avg BS* | 15 | 0.93 | 0.61 | 2681 | 0.55 | 63 | 77 | 0.285 |
| *Avg BS* | 20 | 1.03 | 0.59 | 2683 | 0.56 | 61 | 79 | 0.251 |
| *Avg BS* | 25 | 0.92 | 0.63 | 2643 | 0.55 | 60 | 80 | 0.307 |
| *Avg BS* | 30 | 0.8 | 0.61 | 2645 | 0.55 | 61 | 79 | 0.323 |
| *Avg BS* | 35 | 0.77 | 0.76 | 2489 | 0.51 | 65 | 75 | 0.831 |
| *Avg BS* | 40 | 0.69 | 0.51 | 2607 | 0.54 | 61 | 79 | 0.408 |
| *VulSD* | 0 | 1.04 | 1.05 | 2617 | 0.54 | 67 | 73 | 0.476 |
| *VulSD* | 5 | 1.05 | 1.05 | 2675 | 0.55 | 68 | 72 | 0.345 |
| *VulSD* | 10 | 1.06 | 1.04 | 2700 | 0.55 | 66 | 74 | 0.282 |
| *VulSD* | 15 | 1.06 | 1.01 | 2497 | 0.51 | 63 | 77 | 0.766 |
| *VulSD* | 20 | 1.04 | 1 | 2443 | 0.51 | 61 | 79 | 0.89 |
| *VulSD* | 25 | 1.04 | 1.06 | 2265.5 | 0.47 | 60 | 80 | 0.573 |
| *VulSD* | 30 | 1.05 | 1.1 | 2154 | 0.45 | 61 | 79 | 0.284 |
| *VulSD* | 35 | 1.04 | 1.12 | 2079 | 0.43 | 65 | 75 | 0.135 |
| *VulSD* | 40 | 1.02 | 1.08 | 2071 | 0.43 | 61 | 79 | 0.155 |
| *GenSD* | 0 | 0.9 | 1 | 1590 | 0.33 | 67 | 73 | <0.001 |
| *GenSD* | 5 | 0.91 | 0.99 | 1780 | 0.36 | 68 | 72 | 0.005 |
| *GenSD* | 10 | 0.9 | 0.98 | 1719 | 0.35 | 66 | 74 | 0.003 |
| *GenSD* | 15 | 0.88 | 0.95 | 1832 | 0.38 | 63 | 77 | 0.013 |
| *GenSD* | 20 | 0.93 | 0.96 | 1823 | 0.38 | 61 | 79 | 0.014 |
| *GenSD* | 25 | 0.9 | 0.96 | 1864 | 0.39 | 60 | 80 | 0.024 |
| *GenSD* | 30 | 0.93 | 0.93 | 2052 | 0.43 | 61 | 79 | 0.134 |
| *GenSD* | 35 | 0.9 | 0.94 | 2035 | 0.42 | 65 | 75 | 0.093 |
| *GenSD* | 40 | 0.9 | 0.98 | 1778 | 0.37 | 61 | 79 | 0.008 |
| *MFCL* | 0 | 0.94 | 0.84 | 3053 | 0.62 | 67 | 73 | 0.011 |
| *MFCL* | 5 | 0.92 | 0.81 | 3014.5 | 0.62 | 68 | 72 | 0.018 |
| *MFCL* | 10 | 0.92 | 0.86 | 2950 | 0.6 | 66 | 74 | 0.034 |
| *MFCL* | 15 | 0.9 | 0.87 | 2741 | 0.57 | 63 | 77 | 0.187 |
| *MFCL* | 20 | 0.92 | 0.83 | 2926 | 0.61 | 61 | 79 | 0.03 |
| *MFCL* | 25 | 0.93 | 0.81 | 3173 | 0.66 | 60 | 80 | 0.001 |
| *MFCL* | 30 | 0.92 | 0.79 | 3058 | 0.63 | 61 | 79 | 0.006 |
| *MFCL* | 35 | 0.85 | 0.79 | 2911 | 0.6 | 65 | 75 | 0.048 |
| *MFCL* | 40 | 0.85 | 0.77 | 3133.5 | 0.65 | 61 | 79 | 0.002 |
| *AvgPPMR* | 0 | 0.93 | 0.97 | 2209 | 0.45 | 67 | 73 | 0.325 |
| *AvgPPMR* | 5 | 0.9 | 0.98 | 2168 | 0.44 | 68 | 72 | 0.244 |
| *AvgPPMR* | 10 | 0.89 | 0.96 | 2145 | 0.44 | 66 | 74 | 0.216 |
| *AvgPPMR* | 15 | 0.88 | 0.96 | 2103 | 0.43 | 63 | 77 | 0.177 |
| *AvgPPMR* | 20 | 0.87 | 0.96 | 1982 | 0.41 | 61 | 79 | 0.073 |
| *AvgPPMR* | 25 | 0.88 | 0.99 | 1707 | 0.36 | 60 | 80 | 0.004 |
| *AvgPPMR* | 30 | 0.88 | 0.99 | 1862 | 0.39 | 61 | 79 | 0.022 |
| *AvgPPMR* | 35 | 0.93 | 1.02 | 1817 | 0.37 | 65 | 75 | 0.01 |
| *AvgPPMR* | 40 | 0.91 | 1.04 | 1515 | 0.31 | 61 | 79 | <0.001 |
| *B* | 0 | 0.97 | 1.07 | 1748.5 | 0.36 | 67 | 73 | 0.004 |
| *B* | 5 | 0.96 | 1.07 | 1904.5 | 0.39 | 68 | 72 | 0.024 |
| *B* | 10 | 0.93 | 1.09 | 1662.5 | 0.34 | 66 | 74 | 0.001 |
| *B* | 15 | 0.92 | 1.04 | 1790 | 0.37 | 63 | 77 | 0.008 |
| *B* | 20 | 0.93 | 1.06 | 1711.5 | 0.36 | 61 | 79 | 0.003 |
| *B* | 25 | 0.9 | 1.1 | 1496.5 | 0.31 | 60 | 80 | <0.001 |
| *B* | 30 | 0.93 | 1.1 | 1702.5 | 0.35 | 61 | 79 | 0.003 |
| *B* | 35 | 0.92 | 1.1 | 1745 | 0.36 | 65 | 75 | 0.004 |
| *B* | 40 | 0.94 | 1.14 | 1464.5 | 0.3 | 61 | 79 | <0.001 |
| *I* | 0 | 0.98 | 0.89 | 3131.5 | 0.64 | 67 | 73 | 0.004 |
| *I* | 5 | 0.99 | 0.86 | 3200 | 0.65 | 68 | 72 | 0.002 |
| *I* | 10 | 0.99 | 0.78 | 3200 | 0.66 | 66 | 74 | 0.002 |
| *I* | 15 | 0.97 | 0.89 | 2973.5 | 0.61 | 63 | 77 | 0.022 |
| *I* | 20 | 1 | 0.82 | 3036 | 0.63 | 61 | 79 | 0.008 |
| *I* | 25 | 1.02 | 0.77 | 3234 | 0.67 | 60 | 80 | <0.001 |
| *I* | 30 | 0.95 | 0.79 | 3056.5 | 0.63 | 61 | 79 | 0.006 |
| *I* | 35 | 0.94 | 0.76 | 3046 | 0.62 | 65 | 75 | 0.011 |
| *I* | 40 | 0.94 | 0.71 | 3120 | 0.65 | 61 | 79 | 0.003 |
| *T* | 0 | 1.11 | 1.27 | 2351 | 0.48 | 67 | 73 | 0.695 |
| *T* | 5 | 1.14 | 1.26 | 2305 | 0.47 | 68 | 72 | 0.552 |
| *T* | 10 | 1.21 | 1.3 | 2285 | 0.47 | 66 | 74 | 0.514 |
| *T* | 15 | 1.26 | 1.27 | 2477 | 0.51 | 63 | 77 | 0.831 |
| *T* | 20 | 1.25 | 1.27 | 2393 | 0.5 | 61 | 79 | 0.946 |
| *T* | 25 | 1.28 | 1.27 | 2346.5 | 0.49 | 60 | 80 | 0.823 |
| *T* | 30 | 1.33 | 1.31 | 2392.5 | 0.5 | 61 | 79 | 0.945 |
| *T* | 35 | 1.33 | 1.35 | 2367 | 0.49 | 65 | 75 | 0.77 |
| *T* | 40 | 1.41 | 1.38 | 2309.5 | 0.48 | 61 | 79 | 0.676 |
| *Q* | 0 | 0.93 | 0.9 | 2421 | 0.49 | 67 | 73 | 0.92 |
| *Q* | 5 | 0.93 | 0.87 | 2771 | 0.57 | 68 | 72 | 0.179 |
| *Q* | 10 | 0.94 | 0.9 | 2728 | 0.56 | 66 | 74 | 0.233 |
| *Q* | 15 | 0.89 | 0.86 | 2742 | 0.57 | 63 | 77 | 0.186 |
| *Q* | 20 | 0.87 | 0.85 | 2592 | 0.54 | 61 | 79 | 0.444 |
| *Q* | 25 | 0.85 | 0.86 | 2524 | 0.53 | 60 | 80 | 0.603 |
| *Q* | 30 | 0.83 | 0.86 | 2535 | 0.53 | 61 | 79 | 0.599 |
| *Q* | 35 | 0.83 | 0.89 | 2440 | 0.5 | 65 | 75 | 0.993 |
| *Q* | 40 | 0.86 | 0.86 | 2387.5 | 0.5 | 61 | 79 | 0.928 |
| *MxSim* | 0 | 0.93 | 0.9 | 2421 | 0.49 | 67 | 73 | 0.92 |
| *MxSim* | 5 | 0.7 | 0.74 | 2462 | 0.5 | 67 | 73 | 0.947 |
| *MxSim* | 10 | 0.72 | 0.74 | 2455 | 0.5 | 68 | 72 | 0.978 |
| *MxSim* | 15 | 0.67 | 0.66 | 2582 | 0.53 | 66 | 74 | 0.56 |
| *MxSim* | 20 | 0.65 | 0.68 | 2327 | 0.48 | 63 | 77 | 0.681 |
| *MxSim* | 25 | 0.72 | 0.67 | 2496 | 0.52 | 61 | 79 | 0.718 |
| *MxSim* | 30 | 0.73 | 0.66 | 2769 | 0.58 | 60 | 80 | 0.121 |
| *MxSim* | 35 | 0.69 | 0.64 | 2588 | 0.54 | 61 | 79 | 0.454 |
| *MxSim* | 40 | 0.67 | 0.65 | 2524 | 0.52 | 65 | 75 | 0.719 |
| *Total Biomass* | 0 | 0.49 | 0.45 | 2673 | 0.55 | 67 | 73 | 0.344 |
| *Total Biomass* | 5 | 0.44 | 0.45 | 2462 | 0.5 | 68 | 72 | 0.955 |
| *Total Biomass* | 10 | 0.56 | 0.45 | 2469 | 0.51 | 66 | 74 | 0.912 |
| *Total Biomass* | 15 | 0.62 | 0.42 | 2666 | 0.55 | 63 | 77 | 0.315 |
| *Total Biomass* | 20 | 0.65 | 0.47 | 2587 | 0.54 | 61 | 79 | 0.457 |
| *Total Biomass* | 25 | 0.69 | 0.5 | 2600 | 0.54 | 60 | 80 | 0.401 |
| *Total Biomass* | 30 | 0.75 | 0.58 | 2517 | 0.52 | 61 | 79 | 0.653 |
| *Total Biomass* | 35 | 0.8 | 0.66 | 2520 | 0.52 | 65 | 75 | 0.732 |
| *Total Biomass* | 40 | 0.81 | 0.8 | 2424 | 0.5 | 61 | 79 | 0.953 |
| *community stability* | 0 | 0.79 | 0.66 | 2645 | 0.54 | 67 | 73 | 0.406 |
| *community stability* | 5 | 0.6 | 0.59 | 2443 | 0.5 | 68 | 72 | 0.985 |
| *community stability* | 10 | 0.32 | 0.44 | 2083.5 | 0.43 | 66 | 74 | 0.132 |
| *community stability* | 15 | 0.25 | 0.29 | 2331 | 0.48 | 63 | 77 | 0.688 |
| *community stability* | 20 | 0 | 0 | 2248 | 0.47 | 61 | 79 | 0.462 |
| *community stability* | 25 | 0 | 0 | 2286 | 0.48 | 60 | 80 | 0.552 |
| *community stability* | 30 | 0 | 0 | 2388 | 0.5 | 61 | 79 | 0.908 |
| *community stability* | 35 | 0 | 0 | 2210 | 0.45 | 65 | 75 | 0.103 |
| *community stability* | 40 | 0 | 0 | 2486 | 0.52 | 61 | 79 | 0.621 |
| *population stability* | 0 | 0.88 | 0.95 | 2329 | 0.48 | 67 | 73 | 0.628 |
| *population stability* | 5 | 0.75 | 0.84 | 2408 | 0.49 | 68 | 72 | 0.869 |
| *population stability* | 10 | 0.41 | 0.76 | 2028.5 | 0.42 | 66 | 74 | 0.083 |
| *population stability* | 15 | 0.19 | 0.44 | 2209.5 | 0.46 | 63 | 77 | 0.362 |
| *population stability* | 20 | 0.1 | 0.18 | 2191 | 0.45 | 61 | 79 | 0.342 |
| *population stability* | 25 | 0.02 | 0.08 | 2360 | 0.49 | 60 | 80 | 0.861 |
| *population stability* | 30 | 0.13 | 0.11 | 2615 | 0.54 | 61 | 79 | 0.384 |
| *population stability* | 35 | 0.12 | 0.11 | 2632 | 0.54 | 65 | 75 | 0.414 |
| *population stability* | 40 | 0.12 | 0.11 | 2405 | 0.5 | 61 | 79 | 0.986 |

**Figure S1. *Effect of temperature on (a) Species richness, (b) connectance, (c) mean food chain length, (d) vulnerability, (e) modularity and (f) generality.*** Plots show the mean (points) and standard deviation (bars) across 140 replicated simulations per temperature value from 0 to 40°C, every 1°C. The same 140 distinct food webs were used across the 41 temperature regimes. See *Material and Methods* on the main text for details.

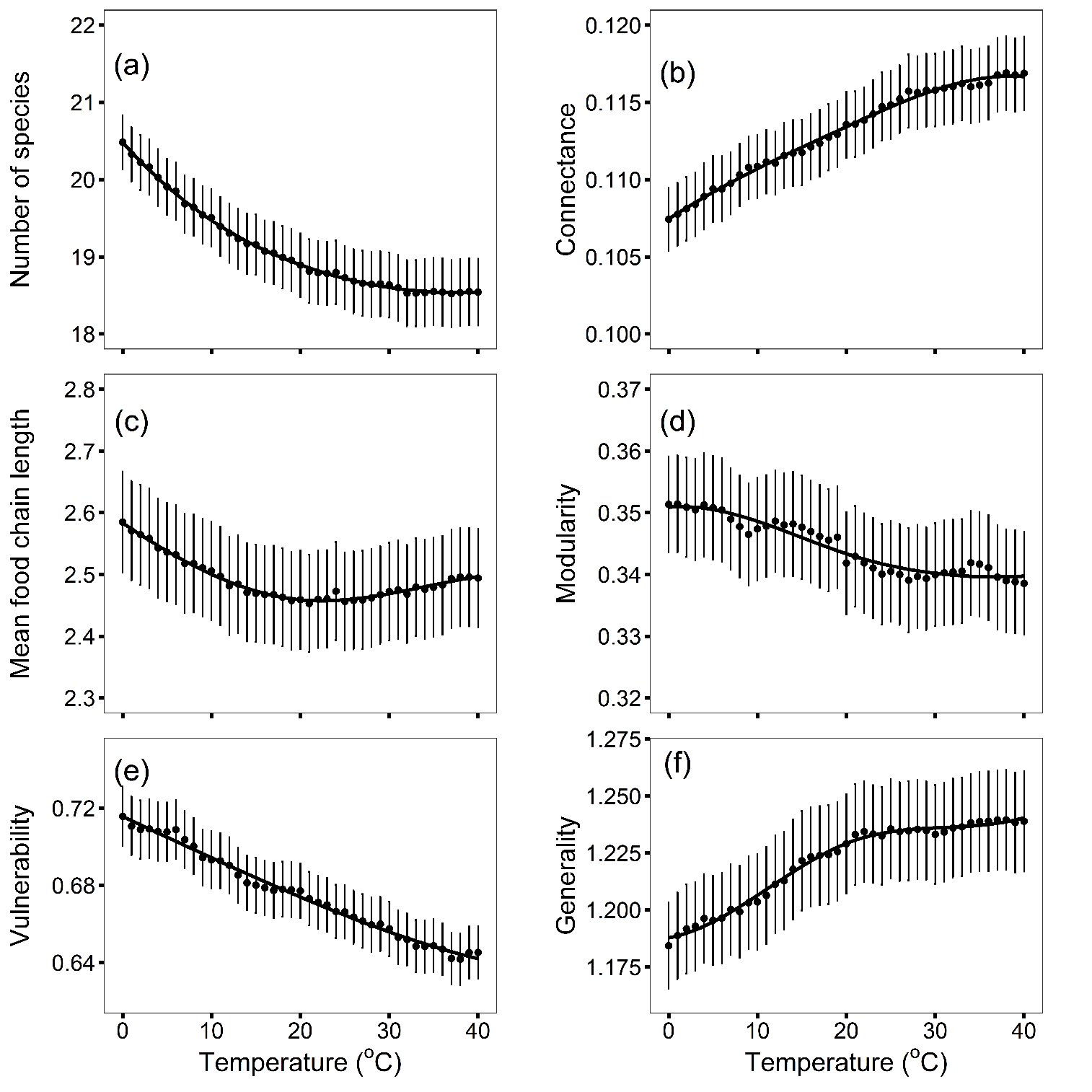

**Figure S2.** ***Proportion of top (a), intermediate (b) and basal (c) species within food webs before invasion along the temperature gradient.*** Plots show the mean (points) and standard deviation (bars) across 140 replicated simulations per temperature value from 0 to 40°C, every 1°C. The same 140 distinct food webs were used across the 41 temperature regimes. See *Material and Methods* on the main text for details.

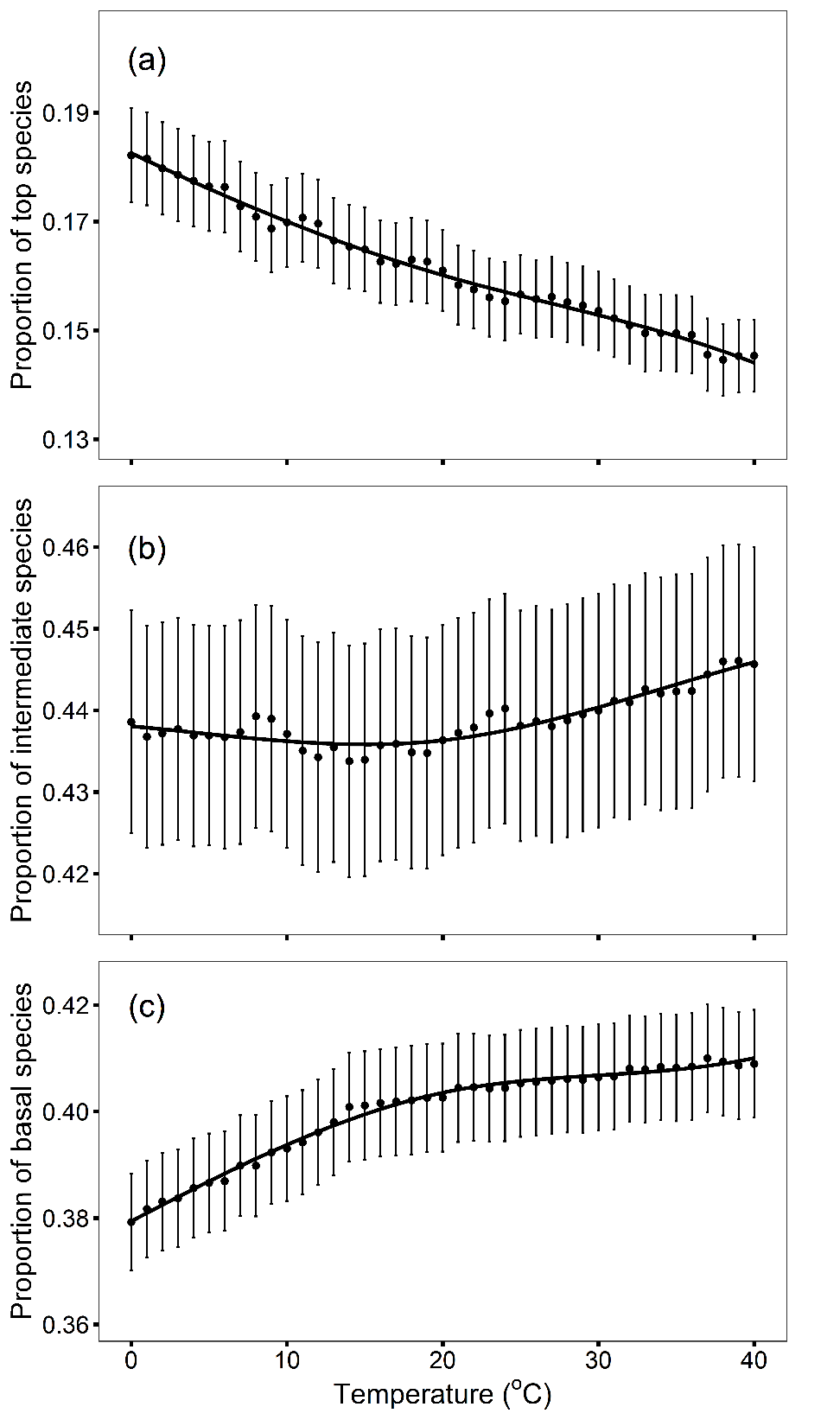

**Figure S3. *Total community biomass (a), community level biomass stability (b) and population level biomass stability (c) as a function of temperature.*** Plots show the mean (points) and standard deviation (bars) across 140 replicated simulations per temperature value from 0 to 40°C, every 1°C. The same 140 distinct food webs were used across the 41 temperature regimes. See *Material and Methods* on the main text for details.

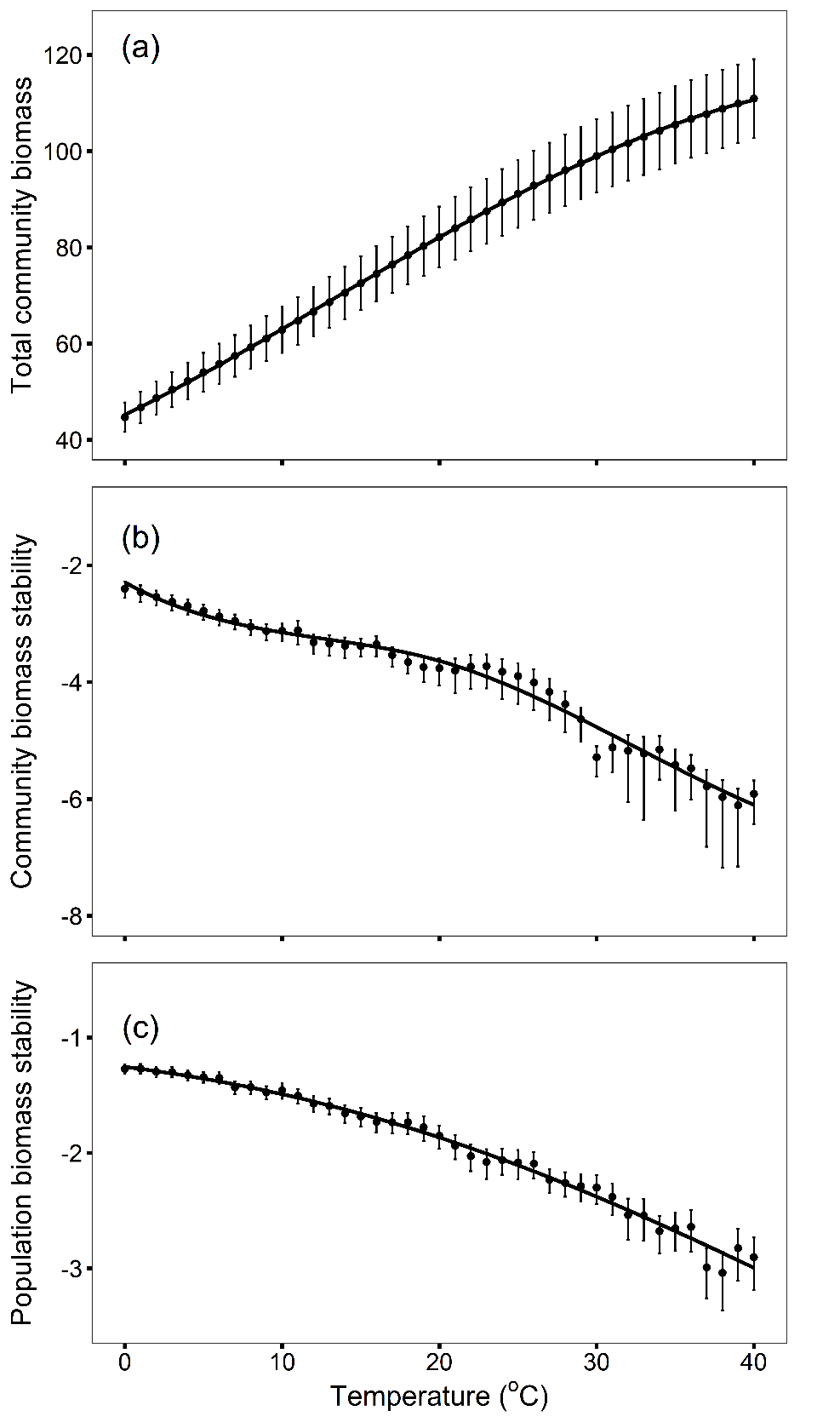

**Figure S4. *Effect sizes (mean ± s.d.) of unsuccessful invasions on complex food webs across temperature regimes on network (a) and community (b) properties.*** Effect sizes were quantified as the ratio between the value of each property after species introduction vs. before introduction, minus unity. Negative values thus indicate negative effects of the invasion on the community (i.e. the value after the invader’s establishment is smaller than before the introduction). Only effects on network/community properties that were found to be significantly different than those observed in successful invasions (Fig. 2 in main text) are shown (Table S3).

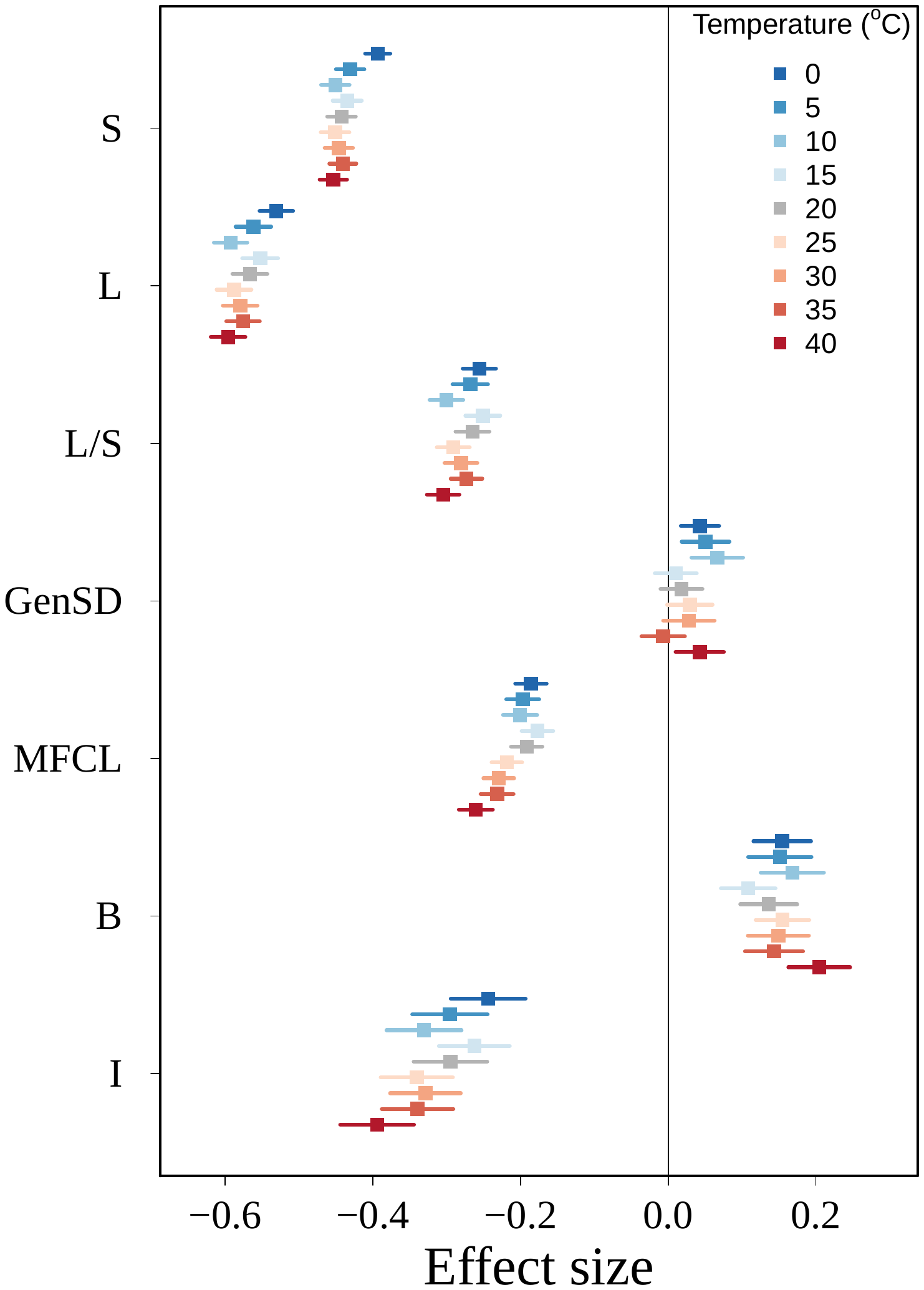
